# A drug’s most potent target is not necessarily the source of its anti-cancer activity

**DOI:** 10.1101/2022.10.16.512438

**Authors:** Debanjan Bhattacharjee, Jaweria Bakar, Erin L. Sausville, Brianna E. Mendelson, Kaitlin Long, Joan C. Smith, Jason M. Sheltzer

## Abstract

The small-molecule drug ralimetinib was developed as an inhibitor of the kinase p38α, and it has advanced to phase 2 clinical trials in oncology. Here, we apply a multi-modal approach to demonstrate that ralimetinib’s anti-cancer activity occurs due to its ability to inhibit EGFR, rather than p38α. We find that cancer cell lines driven by EGFR mutations exhibit the greatest sensitivity to ralimetinib treatment, and ralimetinib phenocopies established EGFR inhibitors in pharmacogenomic profiling experiments. We further demonstrate that ralimetinib inhibits EGFR kinase activity *in vitro* and *in cellulo*, albeit at >30-fold higher concentrations than it inhibits p38α. Finally, while deletion of the gene encoding p38α has no effect on ralimetinib sensitivity, expression of the EGFR-T790M gatekeeper mutation confers resistance to ralimetinib treatment. These findings suggest that future clinical trials involving ralimetinib could incorporate EGFR mutation status as a biomarker to identify sensitive patients. Moreover, our results demonstrate that a compound’s anti-cancer effects should not necessarily be attributed to the protein that it inhibits most strongly, and instead, comprehensive cellular and genetic profiling is required to understand a drug’s mechanism-of-action.

## INTRODUCTION

Despite the enormous potential offered by cancer precision medicine, 97 out of every 100 drug-indication pairs that are tested in clinical trials in oncology fail to receive FDA approval^1^. Retrospective analyses indicate that many drugs fail to progress because their administration results in significant toxicity or has little effect on disease progression^2–4^. It is at present unclear why so many new drugs encounter these problems. Recently, our lab and several other investigators have demonstrated that many drugs that enter oncology clinical trials do so with an incorrect understanding of their mechanism-of-action (MOA)^5–12^. That is, a drug may be developed as an inhibitor of a specific protein, but careful genetic experiments reveal that its anti-cancer effects do not result from the inhibition of its designated target. Mischaracterization of drug MOAs undermines the potential of cancer precision medicine, as clinical trials may be designed to test these drugs in patients with an irrelevant biomarker or who are otherwise unlikely to exhibit a significant response.

In general, the gold standard for proving a drug’s MOA is by identifying a mutation in the drug’s putative target that causes resistance to its effects^13–15^. However, this level of genetic evidence in support of an MOA is rare. In the absence of a resistance-granting mutation, no uniform criteria exist for assigning a particular MOA to a cancer drug. Commonly, a compound is tested against a panel of enzymes *in vitro*, and then that compound is described as an inhibitor of the enzyme for which it exhibits the lowest IC50 value. In an attempt to establish standard benchmarks for assessing compound MOAs, groups like the Structural Genomics Consortium and the Chemical Probes Portal have published guidelines for evaluating drug specificity. These criteria include requirements that the drug has an IC50 against its target that is less than 100 nM and that it exhibits >30-fold selectivity over other enzymes in its target family^16,17^. Beyond *in vitro* enzymatic profiling, several methods have been developed to identify the MOAs of poorly characterized compounds (reviewed in ref^18^). Notably, recent large-scale drug-perturbation assays have facilitated pharmacogenomic-based approaches for drug target identification^19–22^: by assessing cellular phenotypes like gene expression, viability, or morphology in the presence of different chemical perturbations, drugs with unknown MOAs can be linked by similarity to specific pathways or to other drugs with established MOAs.

Uncovering the true MOA of a mischaracterized drug may shed light on its performance in the clinic and could lead to beneficial repurposing studies in alternate cancer types. Toward that end, in this work we describe our study of the small-molecule compound ralimetinib (LY2228820). Ralimetinib was developed by Eli Lilly for use in cancer as an inhibitor of the p38α (MAPK14) kinase^23^. However, we previously reported that CRISPR-induced deletions of MAPK14 fail to alter cancer cell sensitivity to ralimetinib, suggesting that its anti-cancer activity occurs independently of p38α inhibition^9^. Thus far, ralimetinib has been tested in five different oncology trials, but it has exhibited minimal evidence of clinical efficacy (Table 1). For instance, when ralimetinib was administered as a monotherapy to patients with advanced solid tumors, 0 of 74 patients achieved a partial or complete response^24^, while in a placebo-controlled combination study in ovarian cancer, patients receiving ralimetinib did not exhibit a significant difference in response rate or survival compared to patients receiving the placebo^25^. The lack of observed clinical responses provides further evidence that ralimetinib’s MOA has not been correctly determined.

**Table 1.** Ralimetinib clinical trials.

| NCT Number | Cancer type(s) | Intervention(s) | Results | Citation |
| --- | --- | --- | --- | --- |
| NCT01393990 | Advanced cancers | Ralimetinib dose-escalation study | 0 of 74 patients achieved a partial or complete response | PMID: 26581242 |
| NCT01663857 | Ovarian cancer | All patients receive carboplatin and gemcitabine, patients are randomized to receive placebo or ralimetinib | No significant difference in survival or response rate between ralimetinib and placebo | PMID: 31791552 |
| NCT02364206 | Glioblastoma | Patients received radiotherapy, temozolomide, and ralimetinib | Median progression-free survival was 12.8 months, no placebo arm was available for comparison | PMID: 32976869 |
| NCT02860780 | Advanced cancers | Patients received prexasertib and ralimetinib | 0 of 9 patients achieved a partial or complete response | PMID: 31707688 |
| NCT02322853 | Breast cancer | Patients received ralimetinib and tamoxifen | None published | N/A |

## RESULTS

### Ralimetinib treatment phenocopies the effects of EGFR ablation or inhibition

In order to gain insight into ralimetinib’s MOA, we assessed drug screening data from the Broad Institute’s PRISM project. This dataset is comprised of cell viability measurements for ∼550 cancer cell lines treated with ∼4500 different chemical compounds, including ralimetinib and ∼950 other targeted oncology agents^21^. We calculated pairwise correlations between the vector of cellular sensitivities to ralimetinib and every other drug included in the PRISM screen (Fig. 1A). The drug that exhibited the strongest correlation with ralimetinib was BMS-599626, an EGFR inhibitor in Phase 1 clinical trials (R = 0.39)^26^. Cell lines whose viability decreased upon treatment with ralimetinib also tended to be killed by exposure to BMS-599626, while cell lines unaffected by ralimetinib were also resistant to BMS-599626 (Fig. 1B). We noted that the top 21 drugs exhibiting the strongest correlation with ralimetinib’s sensitivity profile were all known EGFR inhibitors, including the well-validated FDA-approved EGFR inhibitors gefitinib, osimertinib, afatinib, vandetanib, and lapatinib (Fig. 1A and 1C)^27^. These results suggest that ralimetinib functions like an EGFR inhibitor, as ralimetinib exposure closely phenocopies the sensitivity profiles of established EGFR-targeting agents.

**Figure 1.**
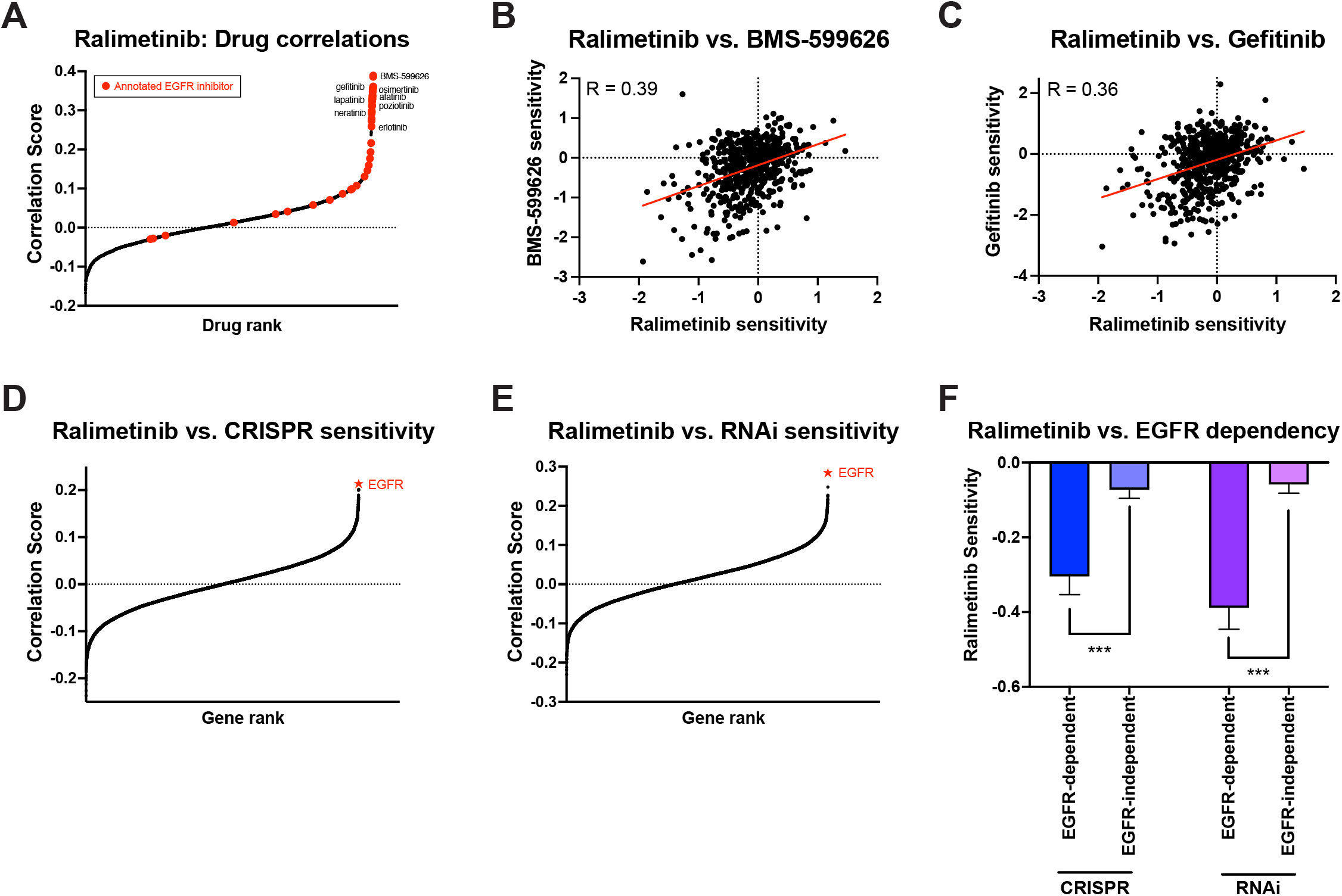
Pharmacogenomic profiling identifies ralimetinib as an EGFR inhibitor. (A) A waterfall plot showing the correlation between the sensitivity profile of ralimetinib and every other drug included in the PRISM dataset^21^. Drugs annotated as EGFR inhibitors are highlighted in red. (B and C) A scatterplot comparing the sensitivity of ∼550 cancer cell lines to ralimetinib and either (B) BMS-599626 or (C) gefitinib. Greater negative scores indicate increasing sensitivity. The red line represents a linear regression plotted against the data. (D and E) A waterfall plot showing the correlation between the sensitivity profile of ralimetinib and the depletion of every gene across a panel of whole-genome (D) CRISPR or (E) RNAi screens^28,29^. The top-scoring gene in both analyses, EGFR, is indicated in red. (F) A bar graph displaying the mean sensitivity of cancer cell lines to ralimetinib, divided based on whether the cell lines require EGFR to proliferate. ***, p < .0001; Student’s t-test.

Next, we sought to determine whether ralimetinib treatment resembles the effects of any specific genetic perturbations. Toward that end, we compared cancer cell line viability following ralimetinib exposure with gene-depletion measurements determined from whole-genome CRISPR and RNAi screens^28,29^. Pairwise correlation analysis revealed that in both the CRISPR and RNAi libraries, the single gene whose ablation most strongly resembled the effects of ralimetinib exposure was EGFR (Fig. 1D-E). Consistent with this finding, we used the CRISPR and RNAi depletion data to divide cancer cell lines into “EGFR-dependent” and “EGFR-independent” subsets, and we observed that ralimetinib treatment resulted in a significantly greater loss of viability among EGFR-dependent cell lines (Fig. 1F). As a comparison, we repeated these analyses with the EGFR inhibitor gefitinib, the BRAF inhibitor vemurafenib, and the PI3K inhibitor alpelisib (Fig. S1). Gefitinib sensitivity resembled ralimetinib and exhibited the strongest correlations with other EGFR-targeting drugs and genetic perturbations (Fig. S1A-C). In contrast, vemurafenib and alpelisib were correlated with BRAF-targeting and PIK3CA-targeting agents, respectively (Fig. S1D-I). These data are consistent with previous reports demonstrating that the targets of anti-cancer agents can be inferred from unbiased pharmacogenomic profiling^12,30^, and they demonstrate that exposure to ralimetinib phenocopies the effects of either genetic or chemical inhibition of EGFR.

### EGFR-mutant cancer cell lines display increased sensitivity to ralimetinib

In order to confirm the results of the PRISM analysis, we assessed the viability of a panel of 20 EGFR-mutant or EGFR-wildtype cancer cell lines treated with ralimetinib. As a control, we exposed these same cell lines to gefitinib in parallel. We performed 7-point dose-response assays to calculate the IC50 values for each drug treatment (Fig. 2A and Table S1). We observed that EGFR-mutant cell lines were significantly more sensitive to ralimetinib compared to cell lines harboring wild-type EGFR (Fig. 2B-2C). The median IC50 value for ralimetinib was 3.6 μM across EGFR-mutant cell lines and 17.2 μM across EGFR-wildtype cell lines. In comparison, ralimetinib was less potent than gefitinib, which exhibited a median IC50 value of 0.02 μM across EGFR-mutant cell lines. However, the overall pattern of cell line sensitivities to ralimetinib and gefitinib was highly-correlated (R = 0.93, P < .0001). These results are consistent with the PRISM analysis and indicate that ralimetinib functions as an EGFR inhibitor.

**Figure 2.**
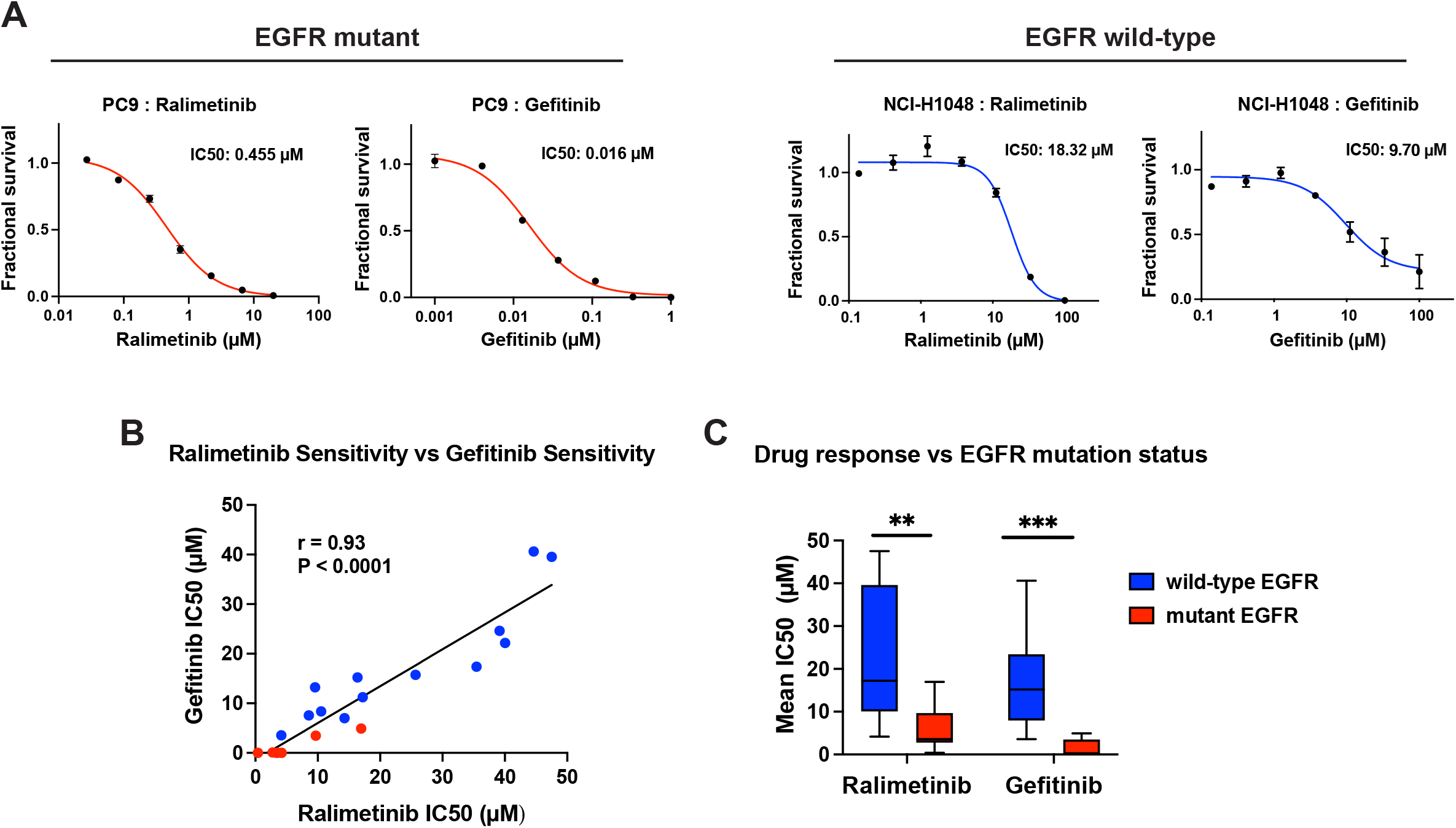
EGFR-mutant cell lines display increased sensitivity to ralimetinib and gefitinib. (A) 7-point dose-response curves displaying cell viability in the presence of the indicated dose of either gefitinib or ralimetinib in either EGFR-mutant (left) or EGFR-wildtype (right) cancer cell lines. (B) A scatterplot showing the correlation between the IC50 values of ralimetinib vs. gefitinib across 20 cancer cell lines. EGFR-mutant cell lines are indicated in red and EGFR-wildtype cell lines are indicated in blue. The black line represents a linear regression plotted against the data. (C) Box plots summarizing the IC50 values of 20 cancer cell lines treated with either ralimetinib or gefitinib, divided based on whether the cell lines harbor an EGFR mutation. **, p < .001; ***, p < .0001; Student’s t-test.

In addition, we noted that two EGFR-mutant cancer cell lines, RL95-2 and NCI-H1975, were relatively resistant to ralimetinib. RL95-2 harbors an A244V mutation in EGFR, but this cell line was resistant to both ralimetinib (IC50 = 9.7 μM) and gefitinib (IC50 = 3.5 μM)^31^. We suspect that this cell line does not depend on EGFR activity to support proliferation. Ralimetinib and gefitinib also displayed reduced potency in NCI-H1975 cells (IC50 = 16.7 μM and 4.9 μM, respectively). This cell line harbors the common L858R activating mutation in EGFR along with a T790M substitution, which causes resistance to first-generation EGFR inhibitors like gefitinib^32^. These results suggest that the T790M mutation blocks the anti-cancer effects of ralimetinib, thereby providing additional evidence that ralimetinib functions as a typical EGFR inhibitor.

### Ralimetinib inhibits EGFR kinase activity *in vitro* and *in cellulo*

As our computational and cellular analyses suggested that ralimetinib functions as an EGFR inhibitor, we performed kinase assays to test whether ralimetinib inhibits EGFR activity *in vitro*. These assays revealed that ralimetinib inhibits the kinase activity of wild-type EGFR (IC50 = 0.175 μM) and several common EGFR-activating mutations (IC50 = 0.013 – 0.232 μM; Fig. 3A). Ralimetinib was potent against EGFR-L858R (IC50 = 0.174 μM), however introduction of the T790M mutation decreased sensitivity to ralimetinib inhibition 20-fold (IC50 = 3.75 μM). Interestingly, among the mutations that we tested, ralimetinib exhibited the highest potency against the relatively-uncommon EGFR-G719C mutation (IC50 = 0.013 μM)^33^. In parallel, we also assessed ralimetinib’s activity against p38α, its previously-reported target, and we observed that it inhibited p38a kinase activity with an IC50 value of 0.004 μM.

**Figure 3.**
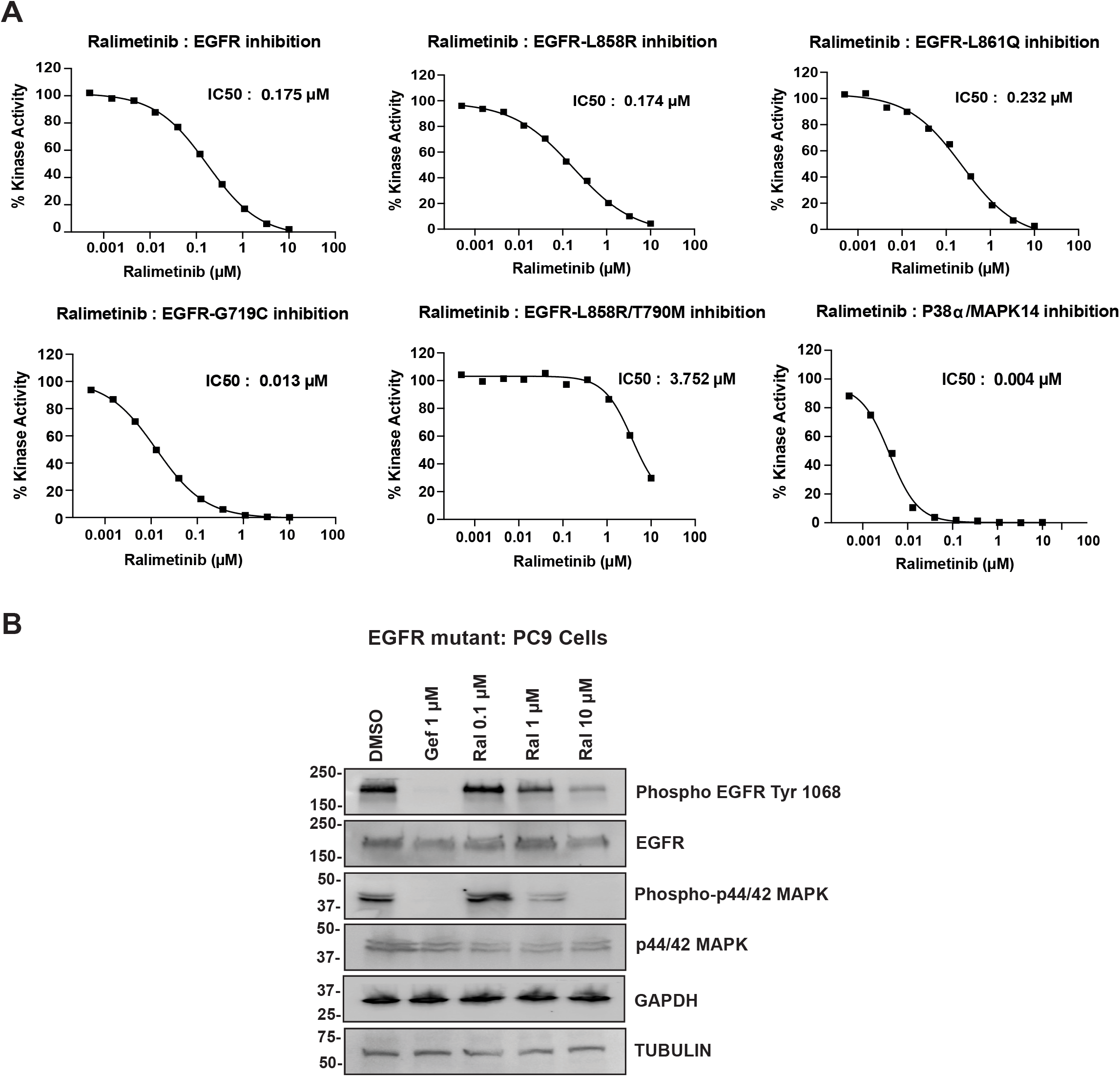
Ralimetinib inhibits EGFR activity *in vitro* and *in cellulo*. (A) Ralimetinib inhibits p38α and various alleles of EGFR in *in vitro* kinase assays. (B) PC9 cells were treated with gefitinib (Gef) or increasing doses of ralimetinib (Ral). After four hours of drug treatment, cell lysates were collected and probed for the phosphorylation of EGFR pathway proteins. Two SDS-PAGE gels were run in parallel and then combined for this panel. One gel was probed for Phospho-EGFR, Phospho-p44/42 MAPK, and Tubulin, and the other one probed for EGFR, p44/42 MAPK, and GAPDH.

In addition, we assessed the effects of ralimetinib on EGFR signaling in EGFR-mutant PC9 cells. We observed that ralimetinib treatment resulted in a dose-dependent decrease in EGFR auto-phosphorylation and phosphorylation of the downstream EGFR target p42/p44-ERK (Fig. 3B). We conclude that ralimetinib functions as an EGFR inhibitor *in vitro* and *in cellulo*.

### The T790M mutation confers resistance to ralimetinib and gefitinib

The gold standard for proving a cancer drug’s mechanism-of-action is the identification of a mutation that grants resistance to its effects^13–15^. We observed that NCI-H1975 cells, which harbor the EGFR-L858R/T790M mutations, were significantly less sensitive to ralimetinib compared to other EGFR-mutant cancer cell lines, and we found that the introduction of T790M decreased the inhibitory effects of ralimetinib in *in vitro* kinase assays (Fig. 2B and 3A). Accordingly, we sought to investigate whether the introduction of this mutation was sufficient to block ralimetinib’s anti-cancer effects. Toward that end, we assessed drug sensitivity in the murine pro-B cell line, Ba/F3. This cell line typically requires exogenous interleukin-3 (IL-3) to support proliferation, but the transgenic expression of a driver oncogene allows the cells to proliferate in the absence of IL-3 and renders them sensitive to small-molecule inhibitors of that oncogene (Fig. 4A)^34^.

**Figure 4.**
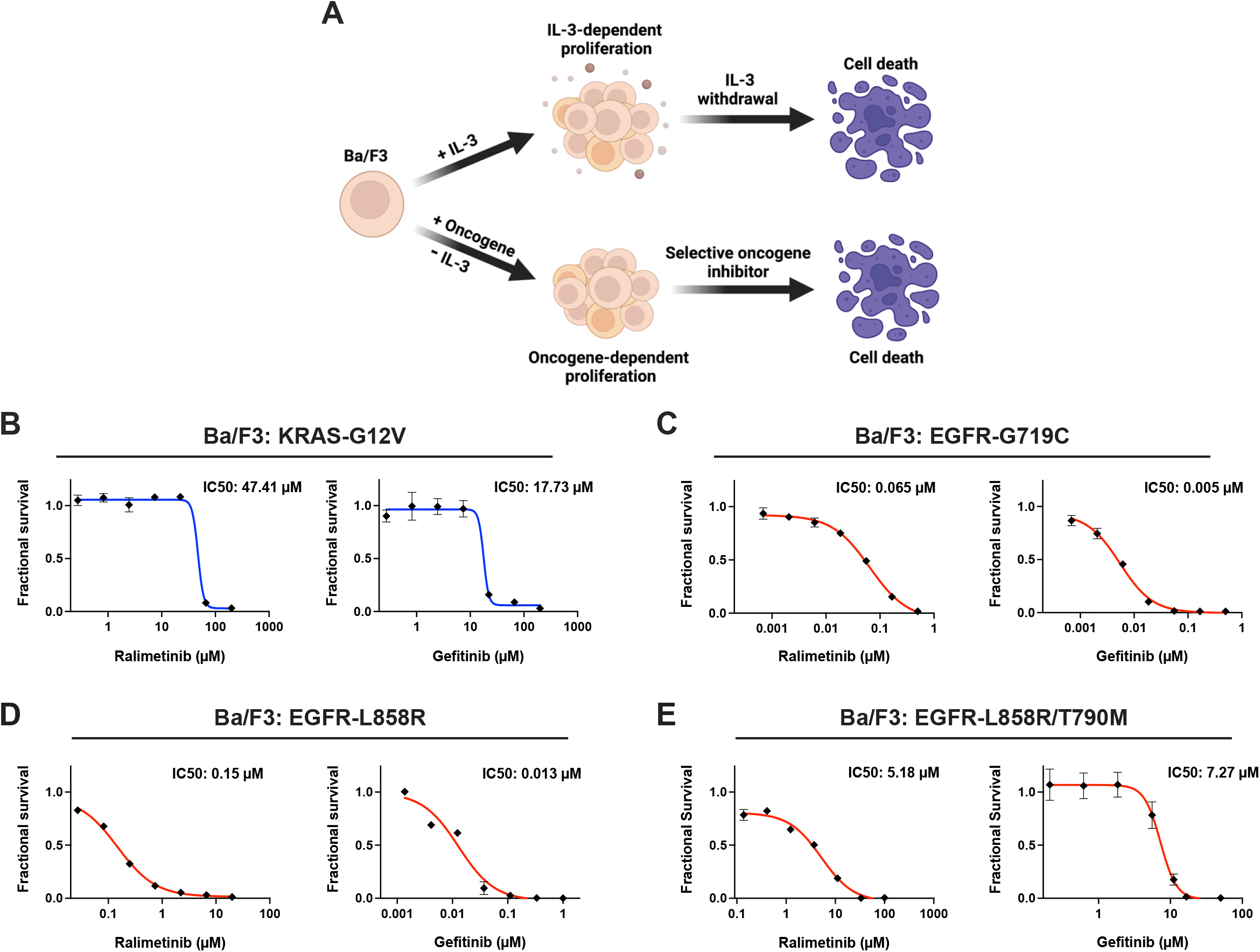
Introduction of the T790M mutation confers resistance to ralimetinib. (A) A schematic of the Ba/F3 system to test oncogene inhibitors. (B-D) 7-point dose-response curves displaying the viability of Ba/F3 cells transduced with the indicated oncogene and then treated with either ralimetinib or gefitinib. Cell lines transduced with a mutant allele of EGFR are indicated in red and cell lines transduced with a different oncogene are indicated in blue.

We transduced Ba/F3 cells with viruses to ectopically express either KRAS-G12V, EGFR-G719C, EGFR-L858R, or EGFR-L858R/T790M, and we observed that all four driver oncogenes supported proliferation in the absence of IL-3. Ba/F3 cells expressing KRAS-G12V displayed minimal sensitivity to either ralimetinib or gefitinib (IC50 = 47.4 and 17.7 μM, respectively; Fig. 4B). In contrast, Ba/F3 cells expressing either EGFR-G719C or EGFR-L858R exhibited sub-micromolar sensitivity to ralimetinib (IC50 = 0.065 and 0.15 μM, respectively; Fig. 4C-D). Finally, introduction of the L858R/T790M double-mutation caused a 35-fold reduction in sensitivity to ralimetinib, and cells expressing this double mutation were similarly resistant to both ralimetinib and gefitinib (IC50 = 5.2 and 7.3 μM, respectively; Fig. 4D). In total, these results demonstrate that in an isogenic setting, introduction of EGFR-L858R sensitizes cells to ralimetinib while the T790M gatekeeper mutation is sufficient to confer ralimetinib resistance.

## Discussion

Multiple lines of evidence indicate that ralimetinib’s anti-cancer activity results from EGFR inhibition, rather than inhibition of its reported target, p38α:

1. Cancer cells harboring CRISPR-induced deletions of p38α remained sensitive to ralimetinib^9^.
2. In both large-scale pharmacogenomic profiling and an in-house confirmatory study, ralimetinib treatment phenocopied EGFR inhibitors and exhibited the greatest potency in EGFR-driven cancer cell lines.
3. Ralimetinib inhibited EGFR kinase activity *in vitro* and *in cellulo*.
4. In an isogenic model system, manipulating cells to rely on EGFR signaling for survival resulted in sub-micromolar sensitivity to ralimetinib.
5. In an isogenic model system, introduction of the EGFR-T790M mutation was sufficient to induce resistance to ralimetinib.

Our results illustrate how multiple complementary approaches, including CRISPR mutagenesis, pharmacogenomic profiling, and targeted molecular assays, can be used to determine the MOA of an anti-cancer compound. Additionally, these results potentially explain the lack of therapeutic efficacy observed with ralimetinib in clinical testing. EGFR mutation status has not been used as a biomarker in any clinical trial with ralimetinib, and the cancer types that ralimetinib has been tested in – breast, ovarian, and glioblastoma – typically do not respond to EGFR inhibitors (Table 1)^35^. While ralimetinib is less potent than established EGFR inhibitors like gefitinib, it is conceivable that greater clinical responses to ralimetinib would be observed if this drug had been administered to patients with cancers driven by EGFR mutations, including non-small cell lung cancer and colon cancer.

We believe that our findings are broadly relevant for the concept of targeted cancer therapies. Consistent with previous results, we observed that ralimetinib can inhibit p38α^24^. Indeed, ralimetinib meets the Structural Genomics Consortium/Chemical Probes Portal criteria for use as a p38α inhibitor, as it has sub-100 nM potency and it exhibits >30-fold selectivity for p38α relative to EGFR (IC50^p38α^ = 5 nM vs. IC50^EGFR^ = 175 nM). Nonetheless, our results show that ralimetinib’s anti-cancer activity results from its ability to inhibit EGFR, rather than p38α. These results highlight the crucial disconnect between the <u>target(s)</u> of a small-molecule and its <u>mechanism-of-action</u>. A drug may exhibit exquisite potency and selectivity for some cellular protein, but if inhibition of that target does not result in the phenotype of interest (e.g., cancer cell killing), that drug’s MOA may be fully independent of its ability to inhibit its designated target. Consistent with these results, we previously observed that 32 out of 32 tested cancer cell lines tolerated CRISPR-induced ablation of MAPK14 without any loss in fitness, further demonstrating that p38α inhibition has minimal effect on cancer cell growth^9^. Note that these results do not rule out some function for p38α inhibition in a non-cell autonomous process, like blocking angiogenesis or promoting tumor immunosurveillance. Here, we focus solely on investigating ralimetinib’s cell-autonomous effects, and additional studies would be required to assess whether ralimetinib and/or p38α inhibition could influence interactions with the cancer microenvironment or host organism.

Finally, our results illustrate the emerging benefits of pharmacogenomic and cell-based methods to understand drug MOAs. While assessing a compound’s ability to bind or inhibit enzyme panels *in vitro* is an important aspect of drug characterization, this approach is fundamentally limited. Notably, one can only discover a specific target for a compound if that protein is included in the tested panel, and it is difficult to infer *in cellulo* selectivity from *in vitro* IC50 values. In contrast, large-scale cellular perturbation assays allow the assessment of a compound’s phenotypic effects in a more relevant environment in which thousands of different proteins are expressed. By comparing the perturbation signature produced by an unknown compound with signatures derived from highly-specific genetic knockdowns or from well-validated cancer drugs, new or mischaracterized drugs can be linked with specific cellular pathways. We speculate that the widespread adoption of pharmacogenomic profiling assays will help decrease the number of new cancer therapies that enter clinical testing with an incorrect understanding of their MOAs.

## MATERIALS AND METHODS

### Tissue culture

The following human cancer cell lines were obtained from the Sheltzer Lab collection: NCI-H1975, HCC827, HCC4006, NCI-H1048, NCI-H3255, PC9, HCC78, DLD1, HCT116, MDA-MB-231, MiaPaCa-2, PC3, A375, A549, A204 and NCI-H1299. The human cancer cell line 11-18 was a kind gift from Dr. Katerina Politi (Yale). The human cancer cell lines NCI-H1838, NCI-H1568, and RL95-2 were purchased from the American Type Culture Collection (ATCC) (Manassas, VA, USA). The identities of all human cancer cell lines used in this study were confirmed using STR profiling (University of Arizona Genetics Core).

NCI-H1975, HCC827, HCC4006, NCI-H3255, 11-18, PC9, NCI-H1568, NCI-H1838, HCC78, and NCI-H1299 cell lines were cultured in Roswell Park Memorial Institute (RPMI)-1640 medium (Lonza; Cat. No. 12–115F/12) supplemented with 10% Fetal Bovine Serum (FBS) (Sigma-Aldrich; Cat. No. F2442), 2 mM glutamine (Lonza; Cat. No. 17–605F), and 100 U/ml penicillin and streptomycin (Life Technologies; Cat. No. 15140122). DLD1, HCT116, MDA-MB-231, MiaPaCa-2, PC3, and A375 cells were cultured in Dulbecco’s Modified Eagle Medium (DMEM) (Thermo Fisher Scientific; Cat. No. 11995–073) supplemented with 10% FBS, 2 mM glutamine, and 100 U/ml penicillin and streptomycin. RL95-2 cells were cultured in DMEM/F12 (Thermo Fisher Scientific; Cat. No. 11320–033) supplemented with 10% FBS, 0.005 mg/ml insulin (Thermo Fisher Scientific; Cat. No. 12585–014), 2 mM glutamine, and 100 U/ml penicillin and streptomycin. A204 cells were cultured in McCoy’s 5A medium (Life Technologies; Cat. No. 16600–108) supplemented with 10% FBS, 2 mM glutamine, and 100 U/ml penicillin and streptomycin. NCI-1048 cells were cultured in DMEM/F12 supplemented with 5% FBS, 0.005 mg/ml insulin, 4.5 mM glutamine, 10 nM hydrocortisone (STEMCELL Technologies; Cat. No. 07926), 30 nM sodium selenite (Sigma-Aldrich; Cat. No. 10102188), 10 nM β-estradiol (Sigma-Aldrich; Cat. No. 50282), 0.01 mg/ml transferrin human (Sigma-Aldrich; Cat. No. 11096370), and 100 U/ml penicillin and streptomycin. A549 cells were cultured in Ham’s F12 medium (Lonza; Cat. No. 12–615F) supplemented with 10% FBS, 2 mM glutamine, and 100 U/ml penicillin and streptomycin.

The mouse Ba/F3 cell line was a kind gift from Dr. James Fraser (UCSF). Ba/F3 cells were cultured in RPMI-1640 medium supplemented with 10% FBS, 2 mM glutamine, and 100 U/ml penicillin and streptomycin. 10 ng/ml IL-3 (STEMCELL Technologies; Cat. No. 78042) was directly added to the medium to support growth in the absence of an oncogene. Oncogene-expressing Ba/F3 cells were maintained in the same culture media without the addition of IL-3. All cell lines were grown in a humidified environment at 37°C with 5% CO2.

### Drug sensitivity assays

Anti-cancer drugs gefitinib (Selleckchem; Cat. No. S1025), ralimetinib (Selleckchem; Cat. No. S1494) and osimertinib (Selleckchem; Cat. No. S7297) were purchased from Selleck Chemicals (Houston, TX, USA). For adherent cancer cell lines, cells were grown (5,000–20,000/well) in 100 μl media on flat-bottomed 96-well culture plates (CELLTREAT; Cat. No. 229196). Anti-cancer therapeutics were added to the plates 12-24 hours after splitting. The highest drug concentration was added to one row of cells, and then 3 to 10-fold serial dilutions were performed. Cell proliferation was determined 72-120 hours after drug treatment, depending upon the growth rate of each cell line. For the suspension cells, 5,000 cells were plated per well in 50 μl of media in round-bottomed 96-well suspension plates (Corning; Cat. No. 2797). Cells were mixed in a 50:50 ratio with 3-fold serially diluted concentrations of anti-cancer drugs for a total volume of 100 μl. Cell proliferation was determined 72 hours after drug treatment. The fraction of cells surviving was quantified using either a MacsQuant Analyzer 10 (Milltenyi Biotec, Germany) or by the Cytation 5 system (BioTek, Santa Clara, California). Cell viability was measured according to the manufacturer’s instructions for both instruments. Data were normalized to the untreated cells and represented by the mean of three to five independent measurements with a standard error <10%. IC50 values were calculated using a four-parameter inhibition vs. response model in Prism 9 (GraphPad Software, San Diego, CA). 7-point dose-response curves were generated from three to six biological replicates.

### Virus generation and transductions

Plasmid DNA was extracted using the GeneJET Plasmid Maxiprep Kit (ThermoFisher Scientific; Cat. No. K0491) for transfections. For the generation of retroviruses, Plat-E cells were used, whereas HEK293T cells were used for lentivirus generation. Calcium phosphate transfection was used to generate EGFR-G719C (Addgene 116254), EGFR-L858R (Addgene 11012), EGFR-L858R/T790M (Addgene 32073), and K-RAS(V12) (Addgene 9052) viruses as previously described^36,37^. Viruses were harvested 48-72 hours post-transfection and filtered through a 0.45 μm membrane. The viral supernatants were immediately added to Ba/F3 cells using 8-10 μg/ml polybrene for transduction. The transduced cells were selected with 0.5 μg/ml puromycin. The transduced Ba/F3 cells were subjected to IL-3 withdrawal after one week or two passages following puromycin selection. The IL-3 independent growth of the transduced cells was monitored for 21 days. This workflow was repeated two times for every construct to create a minimum of two biological replicates.

### Recombinant kinase assays

Kinase assays were performed by Reaction Biology (Malvern, PA, USA) as previously described^38^. The following recombinant kinases were tested: EGFR, EGFR-L858R, EGFR-L861Q, EGFR-G719C, EGFR-L858R/T790M, and p38α. Assays were performed with a top concentration of 10 μM using three-fold serial dilutions.

### Western blotting

Cell lysates were collected using radioimmunoprecipitation assay (RIPA) buffer [50 mM Tris, pH 8.0, 150 mM NaCl, 1% IGEPAL, 0.5% sodium deoxycholate, 0.1% sodium dodecyl sulfate, protease inhibitor cocktail (Sigma-Aldrich; Cat. No. 11836170001), and phosphatase inhibitor cocktail (Sigma-Aldrich; Cat. No. 4906845001)]. Whole-cell lysates were quantified using the DC Protein Assay (Bio-Rad; Cat. No. 5000111). Equal amounts of proteins from each sample were denatured and separated in an 8-16% SDS-PAGE gel. The resolved proteins were transferred onto a polyvinylidene difluoride membrane using the Trans-Blot Turbo Transfer System (Bio-Rad). The membranes were stained using 5% Ponceau S solution, washed using 1x TBS-T (10 mM Tris, 150 mM NaCl, 1% Tween-20), and blocked using 5% non-fat dry milk for an hour at room temperature. After blocking, the membranes were incubated in specific antibodies overnight on an orbital shaker. After overnight incubation, the membranes were washed three times using 1x TBS-T and then incubated in secondary antibody solutions for an hour at room temperature. After three washes, proteins were detected using Clarity Max Western ECL Substrate (Bio-Rad; Cat. No. 1705062). The antibodies and dilutions used are as follows: Phospho-EGFR: Cell Signal Cat. No. 3777S, 1:1000 in 5% milk/TBS-T; EGFR: Cell Signal Cat. No. 4267L, 1:750 in 5% milk/TBS-T; Phospho-p44/p42 MAPK (Erk1/2): Cell Signal Cat. No. 4370S, 1:1000 in 5% milk/TBS-T; p44/p42 MAPK (Erk1/2): Cell Signal Cat. No. 4695S, 1:500 in 5% milk/TBS-T; GAPDH-HRP: Santa Cruz Cat. No sc-365062, 1:1500 in 5% milk/TBS-T; Tubulin: Sigma-Aldrich Cat. No. T6199, 1:5000 in 5% milk/TBS-T; Goat Anti-Rabbit: Abcam Cat. No. 97051, 1:5000 or 1:10000 in 5% milk/TBS-T; Goat Anti-Mouse: Bio-Rad Cat. No. 1706516, 1:10000 in 5% milk/TBS-T.

### Analysis of pharmacogenomic profiling data

Drug, CRISPR, and RNAi screening data were downloaded from www.depmap.org^21,28^. For the drug-drug analysis, pairwise Pearson correlations were calculated between the vector of cellular sensitivities to ralimetinib from the PRISM primary screen and the sensitivity values for every other drug included in the PRISM primary screen. For the drug-gene analyses, pairwise Pearson correlations were calculated between the same ralimetinib values and the gene effect scores from CRISPR or RNAi screening. For the RNAi analysis, the data were preprocessed to include only targets with fewer than 250 missing values. Code for this portion of the analysis is available at https://github.com/sheltzer-lab/ralimetinib.

## ACKNOWLEDGMENTS

We are grateful to Dr. Katerina Politi (Yale University) for providing the 11-18 cell line and to Dr. James Fraser (UCSF) for providing the Ba/F3 cell line.

Research in the Sheltzer Lab is supported by NIH grant R01CA237652, Department of Defense grant W81XWH-20-1-068, an American Cancer Society Research Scholar Grant, a sponsored research agreement from Ono Pharmaceuticals, and a sponsored research agreement from Meliora Therapeutics.

## Supplemental Figure Legends

**Figure S1.**
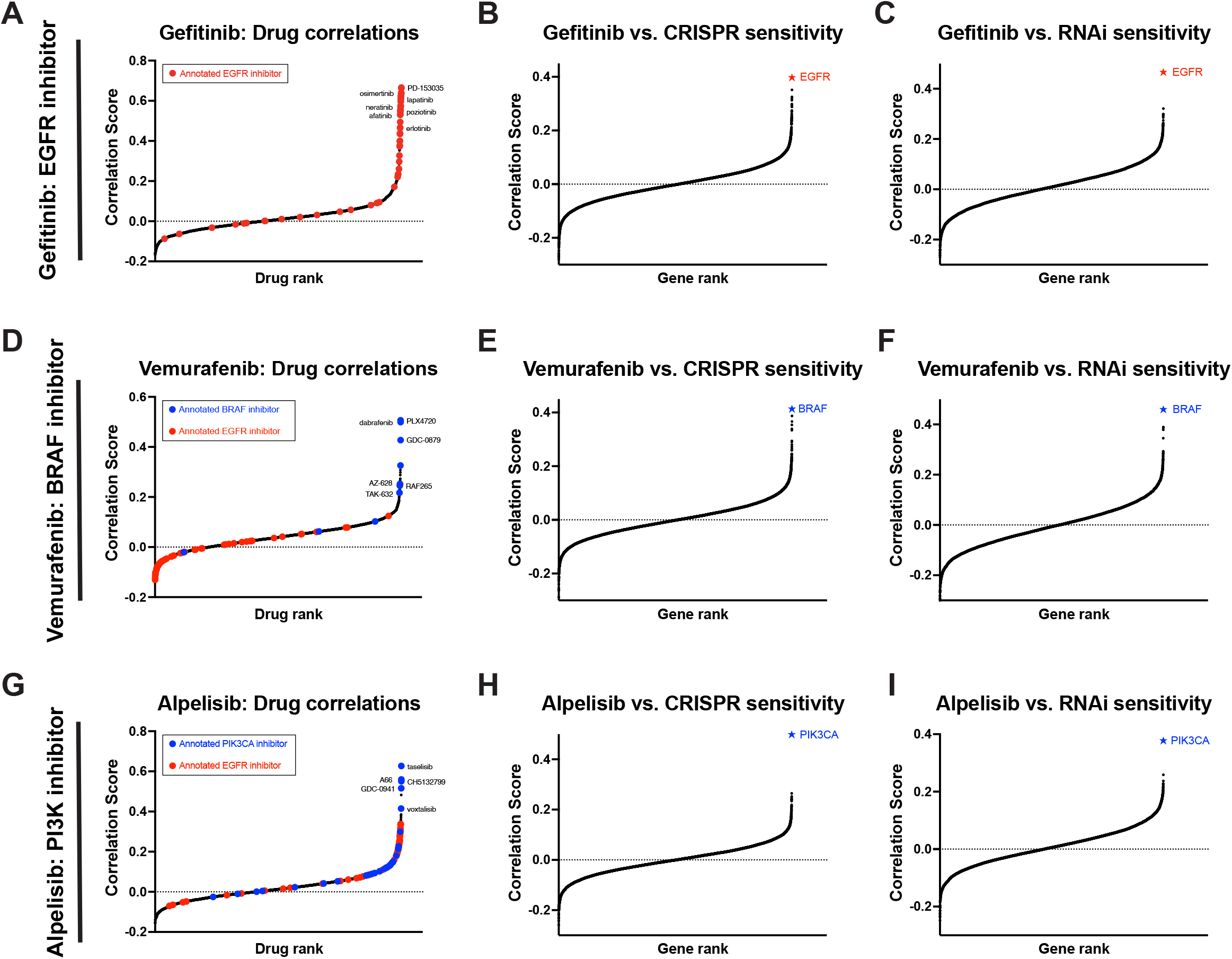
Pharmacogenomic profiling of established EGFR, BRAF, and PIK3CA inhibitors. (A) A waterfall plot showing the correlation between the sensitivity profile of the EGFR inhibitor gefitinib and every other drug included in the PRISM dataset^21^. Drugs annotated as EGFR inhibitors are highlighted in red. (B and C) A waterfall plot showing the correlation between the sensitivity profile of gefitinib and the depletion of every gene across a panel of whole-genome (B) CRISPR or (C) RNAi screens^28,29^. The top-scoring gene in both analyses, EGFR, is indicated in red. (D) A waterfall plot showing the correlation between the sensitivity profile of the BRAF inhibitor vemurafenib and every other drug included in the PRISM dataset^21^. Drugs annotated as EGFR inhibitors are highlighted in red and drugs annotated as BRAF inhibitors are highlighted in blue. (E and F) A waterfall plot showing the correlation between the sensitivity profile of vemurafenib and the depletion of every gene across a panel of whole-genome (E) CRISPR or (F) RNAi screens^28,29^. The top-scoring gene in both analyses, BRAF, is indicated in blue. (G) A waterfall plot showing the correlation between the sensitivity profile of the PIK3CA inhibitor alpelisib and every other drug included in the PRISM dataset^21^. Drugs annotated as EGFR inhibitors are highlighted in red and drugs annotated as PIK3CA inhibitors are highlighted in blue. (H and I) A waterfall plot showing the correlation between the sensitivity profile of vemurafenib and the depletion of every gene across a panel of whole-genome (H) CRISPR or (I) RNAi screens^28,29^. The top-scoring gene in both analyses, PIK3CA, is indicated in blue.

**Table S1.**
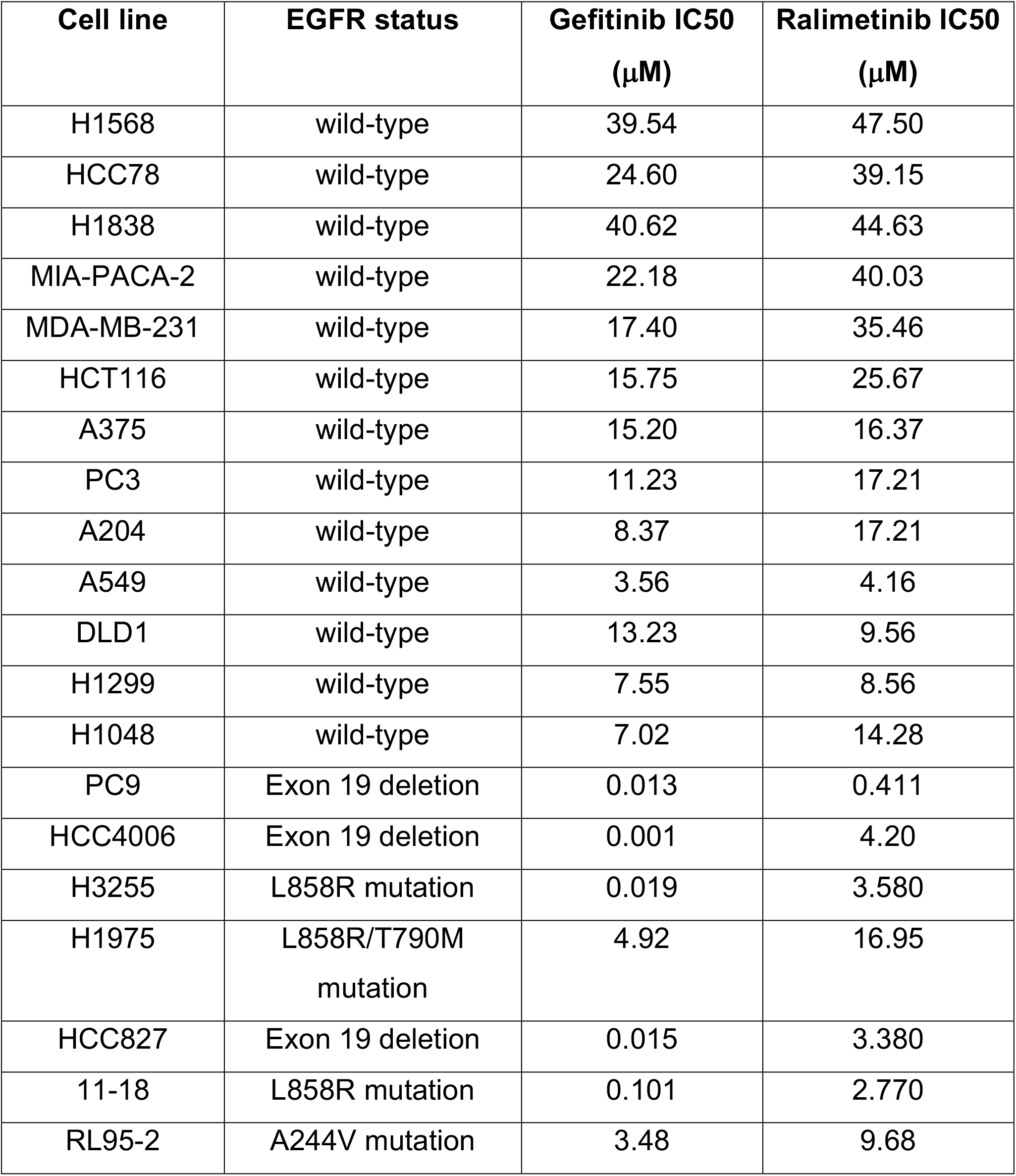
Cellular sensitivity to gefitinib and ralimetinib.

